# Holographic fingerprinting reveals oligomer-driven phase separation in Bovine Serum Albumin

**DOI:** 10.64898/2025.12.10.693513

**Authors:** Fook C. Cheong, Seung Y. Lee, Sujata Bais, Satyam Khanal, Saumya Saurabh

## Abstract

Biomolecular condensates formed through phase separation are fundamental to cellular organization. Although the physical principles underlying intrinsically disordered proteins are well understood, the molecular determinants of condensate formation in globular proteins remain elusive. Here, we employ Holographic Particle Characterization, a label-free, high-throughput imaging technique, to investigate the self-assembly of Bovine Serum Albumin (BSA), a model globular protein. We show that this technique reliably differentiates amorphous aggregates from liquid-like condensates by their distinct refractive indices and morphologies. Coupled with size-exclusion chromatography, our analysis reveals that BSA phase separation strictly depends on higher-order oligomeric assemblies. Monomeric and dimeric fractions fail to form condensates under identical crowding conditions. Furthermore, the internal packing density of these condensates is highly tunable via pH-driven protonation changes but remains insensitive to ionic screening. These findings support a model of “emergent multivalency,” where oligomerization creates a structural scaffold enabling hydrophobically stabilized phase separation, thereby defining a molecular threshold for globular proteins condensation.

## Introduction

Biomolecular condensates formed via phase separation have emerged as a central organizing principle in cell biology, enabling cells to compartmentalize biochemical reactions without membrane boundaries.^1–3^ Within the prevailing stickers-and-spacers framework, multivalent interaction motifs embedded in structured and intrinsically disordered regions collectively drive demixing once a critical saturation concentration is exceeded. ^4–6^ However, many abundant cellular proteins are compact and globular, with limited intrinsic valency and no obvious modular binding domains, raising fundamental questions about how such structured proteins access condensate states in crowded intracellular environments.^7^

Bovine serum albumin (BSA) is a 66.5 kDa globular transport protein and a widely used model for probing phase behavior of folded proteins under macromolecular crowding. Recent studies have shown that BSA can form micrometer-sized, liquid-like droplets in the presence of polyethylene glycol (PEG) and related cosolutes, with phase behavior tuned by protein concentration, temperature, and solution composition. ^8–10^ At the same time, BSA readily undergoes reversible oligomerization into dimers and higher-order assemblies, and can form amorphous aggregates or gels depending on pH, ionic strength, and storage history.^11,12^ This dual propensity for both condensation and aggregation makes BSA an attractive system for disentangling the molecular features that distinguish functional globular-protein condensates from disordered aggregates.^13^

Discriminating between these assembly states remains challenging with standard characterization methods. Bulk turbidity and light-scattering measurements report on overall demixing but cannot reliably differentiate dense liquid droplets from fractal aggregates or quantify their internal packing density.^14^ Fluorescence microscopy, while powerful for visualizing condensates in cells and in vitro, typically relies on extrinsic labels that may perturb surface charge or hydrophobic interactions, potentially altering the very forces that underlie phase separation, especially for charge-sensitive proteins such as BSA.^15,16^ In contrast, holographic particle characterization uses inline digital holograms to extract both radius and refractive index of individual particles.^17^ This approach has been applied for labelfree, high-throughput differentiation of protein aggregates, oil droplets, and other subvisible species based on their optical fingerprints.^18–21^ Recent applications of this technique have demonstrated quantitative sizing and refractive index measurements of protein aggregates and biomolecular condensates, underscoring its potential for dissecting condensate material properties at the single-particle level. ^22,23^

Despite growing evidence that BSA can undergo phase separation under PEG-induced crowding, the molecular prerequisites for this transition remain unresolved. It is not clear whether BSA monomers alone are sufficient to drive condensate formation, or whether specific oligomeric species act as multivalent scaffolds that license phase separation in analogy to multivalent interaction hubs in intrinsically disordered systems.^24^ Likewise, while crowding and depletion interactions are broadly recognized as drivers of phase separation, the precise crowding thresholds and polymer-size requirements that separate reversible condensate formation from irreversible aggregation in globular proteins have not been systematically mapped.^7,25,26^ Furthermore, the relative contributions of electrostatics, hydrophobic interactions, and protonation state in defining the internal density and material properties of BSA condensates remain poorly understood.^9,27–29^

In this work, we use holographic particle characterization to dissect the phase behavior of BSA in PEG-crowded solutions at the single-particle level. By simultaneously measuring size and refractive index for thousands of assemblies, we establish a robust optical criterion that distinguishes high-density liquid condensates from amorphous aggregates and quantify how condensate packing responds to pH, ionic strength, and crowding parameters. Coupling this label-free readout with size-exclusion chromatography, we show that only higher-order oligomeric BSA fractions generate dense condensates under crowding, whereas monomer- and dimer-enriched fractions remain non-condensing or aggregate, supporting a model in which oligomerization provides the emergent multivalency required for globular-protein phase separation. These findings extend multivalency-based theories of phase separation to structured proteins and highlight inline holographic microscopy as a powerful platform for resolving the molecular determinants of condensate formation and maturation.^4,23,24^

## Results and discussion

### Label-Free Differentiation of Protein Aggregates and Condensates

In our previous work, we established holographic particle characterization as a quantitative method to analyze the material properties of condensates formed by the bacterial protein PopZ.^23^ PopZ is an intrinsically disordered protein (IDP) that drives phase separation through the flexible, multivalent interactions typical of low-complexity domains. However, globular proteins like Bovine Serum Albumin (BSA) present a fundamentally different physicochemical challenge. Unlike IDPs, globular proteins possess a defined tertiary structure and inherently lack the extended disorder required to facilitate strong, multivalent interactions. Consequently, distinguishing between their disordered aggregation and controlled phase separation is often difficult using conventional bulk assays. We sought to determine if the holographic “optical fingerprinting” we validated with PopZ could be extended to differentiate the distinct assembly states of a globular protein.

To test this, we induced BSA self-assembly in the presence of a crowding agent (PEG 8K) under two distinct buffer conditions: low ionic strength (1 mM sodium phosphate), known to favor irreversible aggregation, and physiological ionic strength (100 mM sodium phosphate), which supports condensate formation. Inline holographic characterization resolved two distinct BSA assembly states under crowding conditions that favor either aggregation or condensate formation (Figure 1). In the 1 mM buffer, BSA formed particulates exhibiting a broad distribution of refractive indices with a characteristic inverse dependence on size: small particulates displayed high refractive indices, whereas larger assemblies showed progressively lower refractive indices (approaching the buffer index). This trend is indicative of uncontrolled fractal aggregation, where larger clusters become increasingly porous and less dense.^19^ In stark contrast, the 100 mM condition yielded a tight, distinct population of spherical assemblies with consistently high refractive indices (*n*_*p*_ ≈ 1.40–1.44) that were maintained even as the droplets grew in size.^23^

**Figure 1.**
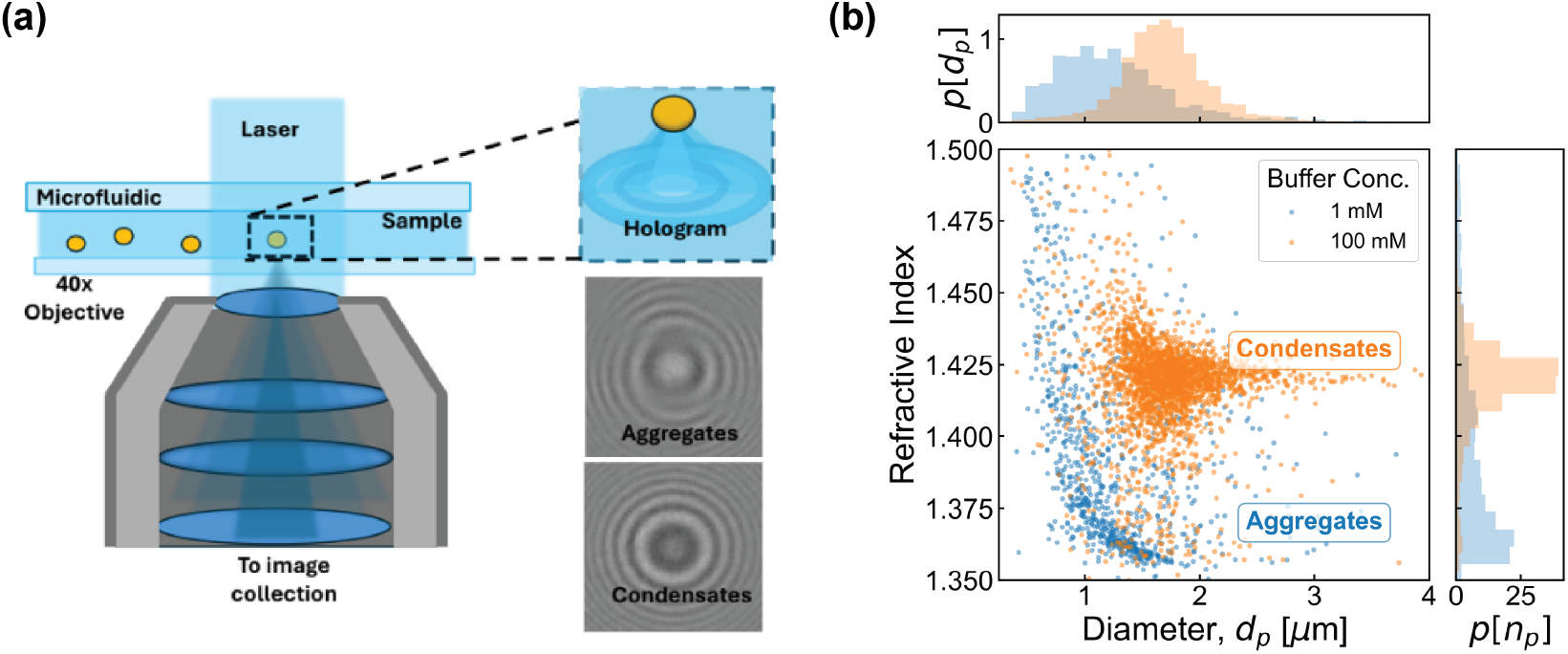
Label-free holographic characterization distinguishes BSA protein aggregates from condensates. (a) Schematic of the inline holographic particle characterization setup. A laser illuminates the sample in a microfluidic channel, and the interference pattern (hologram) created by the unscattered light and the light scattered by a particle is magnified and recorded. Representative holograms are shown for lower-density aggregates and higher-density condensates. (b) A two-dimensional scatter plot of refractive index versus particle diameter (*d*_*p*_) for BSA assemblies. The plot clearly distinguishes two populations based on the buffer conditions: particles formed in 1 mM sodium phosphate buffer (blue) exhibit a broad size distribution with lower refractive indices, characteristic of protein aggregates. In contrast, particles formed in 100 mM sodium phosphate buffer (orange) show a distinct population with significantly higher refractive indices (∼1.40–1.44), corresponding to more densely packed biomolecular condensates.

This single-particle optical fingerprinting directly differentiates condensed liquid droplets from amorphous aggregates without labels or staining, overcoming limitations of turbidity assays and conventional imaging in resolving material state and internal density. Together, these data establish that BSA forms a bona fide high-density liquid phase under appropriate ionic conditions, and that holographic microscopy can robustly isolate this phase for further biophysical analysis.

### Distinct Effects of pH and Ionic Strength on Condensate Density

Having established that BSA condensates represent a distinct, high-density liquid phase, we next investigated the intermolecular forces responsible for maintaining this internal architecture. We asked whether the packing density of these globular protein assemblies could be finely tuned by environmental parameters, or if it represents a fixed material state.

We first probed the effect of pH, which directly alters the protonation states of the protein’s surface residues. Tracking the holographic properties of BSA condensates across a pH range of 4.5 to 7.7 revealed a strong, tunable inverse correlation between pH and condensate density (Figure 2a-b). At acidic conditions (pH 4.5), the protonation of acidic side chains brings the protein close to its isoelectric point (pI ≈ 4.7), effectively neutralizing the net surface charge. We hypothesize that this specific protonation state minimizes intermolecular electrostatic repulsion, allowing the globular domains to achieve closer proximity and higher packing density (*n*_*p*_ ≈ 1.43). As the pH rises towards physiological conditions (pH 7.0–7.7), deprotonation increases the net negative charge on the protein surface, reintroducing repulsive forces that expand the condensate volume and lower the refractive index to approximately 1.405.

**Figure 2.**
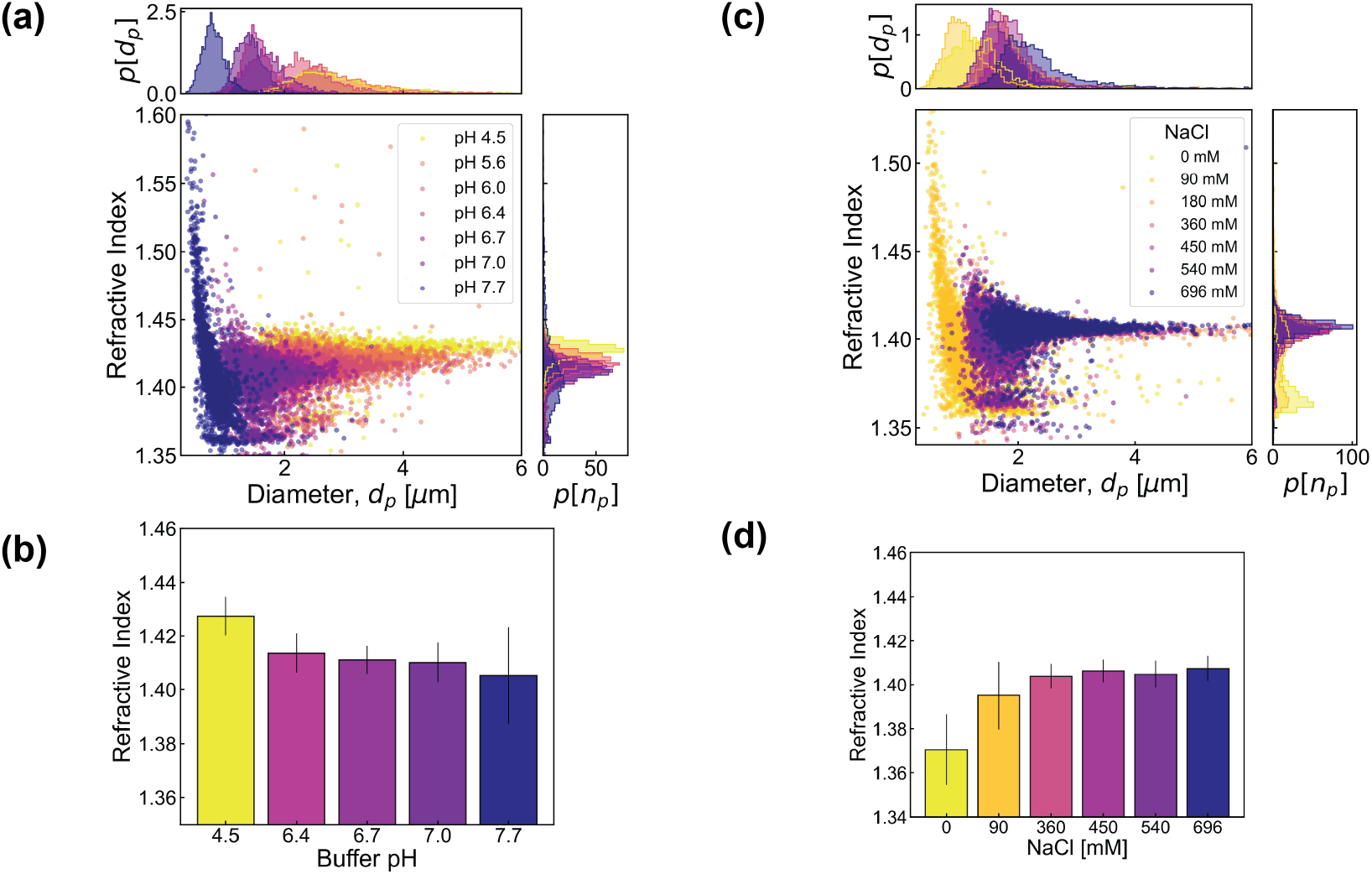
BSA condensate properties are strongly pH-dependent and sensitive within a low ionic strength range. (a) Two-dimensional scatter plots of refractive index versus particle diameter (*d*_*p*_) for BSA condensates formed at varying pH values (pH 4.5 to 7.7). (b) Corresponding bar chart of the mean refractive index as a function of pH. Condensate refractive index shows a strong inverse correlation with pH, decreasing from approximately 1.43 at pH 4.5 to 1.41 at pH 7.03. This trend indicates that more acidic conditions promote larger and more densely packed assemblies. (c) Two-dimensional scatter plots of refractive index versus *d*_*p*_ for condensates formed across a range of NaCl concentrations (0 mM to 696 mM). (d) Corresponding bar chart of the mean refractive index as a function of NaCl concentration. The refractive index remains stable at approximately 1.40–1.41 across all salt concentrations tested. This striking insensitivity to a range of ionic strengths suggests that electrostatic interactions are not the primary drivers for maintaining condensate packing density. All condensates were formed in 100 mM sodium phosphate buffer with 20% (w/v) PEG 8K. Error bars in (b) and (d) represent the standard deviation of the refractive index distribution.

We reasoned that if electrostatic repulsion driven by deprotonation regulates packing density, then screening these charges with ions should similarly modulate the material properties. However, when we varied the sodium chloride concentration from 0 mM to nearly 700 mM, the material properties of the condensates displayed a striking insensitivity to the ionic environment (Figure 2c-d).^28,29^ The refractive index remained constant (*n*_*p*_ ≈ 1.40 − −1.41) across the entire salt range. This robustness demonstrates a fundamental distinction in the mechanism of assembly: while the protonation state of specific residues can *tune* the packing fraction by altering the intrinsic charge landscape, bulk ionic screening cannot. This implies that once the optimal protonation state is established, the cohesive forces maintaining the high-density state are predominantly non-covalent and hydrophobic, rendering the internal architecture resistant to simple Debye screening. With the internal density shown to be robust against ionic screening, we turned our attention to the external forces required to drive the initial phase separation, specifically investigating how the excluded volume and depletion forces imposed by the crowding environment modulate the assembly process.

### Critical Crowding Parameters for Phase Separation

While hydrophobic interactions stabilize the interior of the condensate once formed, BSA is intrinsically soluble and does not phase separate spontaneously in buffer alone. This raises a critical question about the nucleation barrier: what are the specific physical parameters of the crowding environment required to force this globular protein into a dense liquid phase rather than a disordered aggregate?

To define this threshold, we mapped the phase diagram of BSA as a function of the concentration and molecular weight of the crowding agent, polyethylene glycol (PEG) (Figure 3). We first observed a sharp binary transition based on concentration. At PEG 8K concentrations below 20% (w/v), BSA formed only amorphous aggregates. A liquid-liquid phase separation regime emerged exclusively at or above 20% (w/v), where stable, high-refractive-index condensates were observed. Furthermore, within this condensed regime, the refractive index increased monotonically with PEG concentration (Figure 3b). This suggests that the external osmotic pressure exerted by the crowding agent directly compresses the condensate, further confirming that the material state is responsive to depletion forces. ^10,29^ However, excluded volume alone is insufficient to explain this behavior. When we varied the polymer chain length while maintaining constant concentration (20% w/v), we found that molecular weight is a determinant factor (Figure 3c-d). Low molecular weight PEGs (< 6 kDa) failed to induce phase separation, yielding only aggregates, whereas higher molecular weights (> 6 kDa) successfully drove condensate formation. This size-dependence points to a depletion attraction mechanism: the crowding polymer must have a sufficiently large radius of gyration relative to the protein to create an effective depletion zone. Below this threshold, the attractive potential is too weak to sustain the liquid state, and the protein falls into a kinetic trap of aggregation.

**Figure 3.**
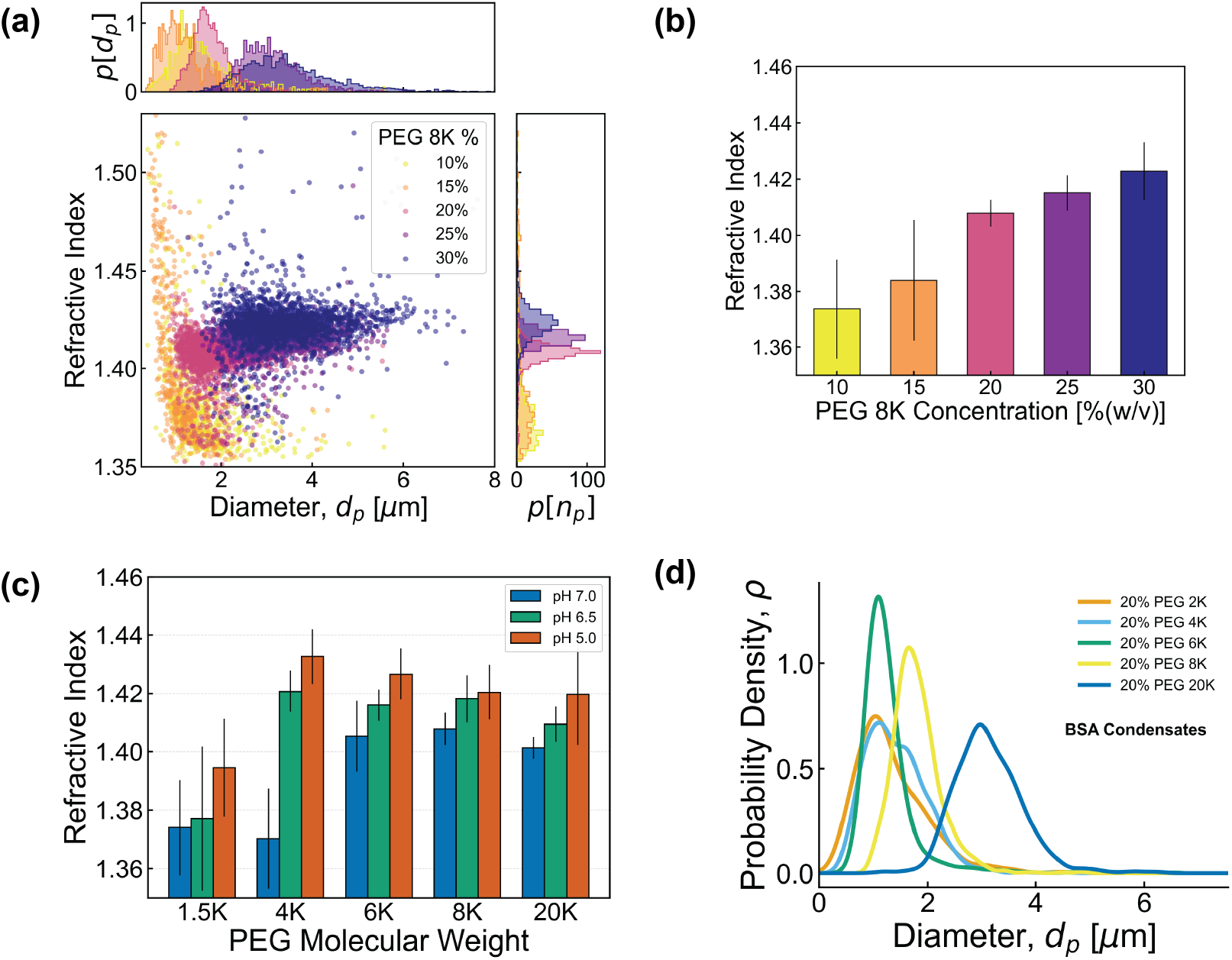
Condensate formation and properties are dependent on the crowding agent. (a) Two-dimensional scatter plots of refractive index versus particle diameter (*d*_*p*_) for BSA assemblies at varying PEG 8K concentrations (10% to 30% w/v). (b) Corresponding bar chart showing the mean refractive index increases with PEG 8K concentration. At concentrations below 20% (w/v), BSA primarily forms large aggregates, while condensates are observed at and above 20% (w/v). The increasing refractive index suggests a more densely packed protein environment as the crowding agent concentration increases. (c) Bar chart of mean refractive index as a function of PEG molecular weight (1.5 KDa to 20 KDa) at pH 5, 6.5, and 7. Condensate formation is dependent on PEG molecular weight; low molecular weight PEGs induce aggregates rather than condensates, while higher molecular weights (>6 KDa) are required to form dense condensates in a pH-dependent manner. Across all molecular weights tested, lower pH conditions consistently produce condensates with higher refractive indices. All experiments were conducted in 100 mM sodium phosphate buffer. Error bars represent the standard deviation of the refractive index distribution.

### Oligomerization as a Prerequisite for Condensate Formation

While depletion forces from crowding agents provide the thermodynamic drive for phase separation, they do not explain the molecular specificity of the process. Why do some globular proteins condense while others aggregate or remain soluble under identical crowding conditions? The current paradigm for protein phase separation relies on “multivalency”— the presence of multiple interaction motifs (“stickers”) connected by flexible linkers. IDPs possess this multivalency intrinsically. Globular proteins like BSA, however, present a compact, solvent-accessible surface that lacks obvious repeating interaction motifs. We hypothe-sized that BSA overcomes this limitation not through conformational disorder, but through *oligomerization*, effectively creating a “multivalent scaffold” from multiple globular subunits. To test this hypothesis, we fractionated BSA using size-exclusion chromatography (SEC) to isolate distinct oligomeric populations (Figure 4a). The elution profile revealed a mixture of monomers, dimers, and higher-order oligomers (likely hexamers and octamers). We collected individual fractions across this range and subjected them to the exact same crowding conditions established in the previous section (20% PEG 8K, 100 mM NaP).

**Figure 4.**
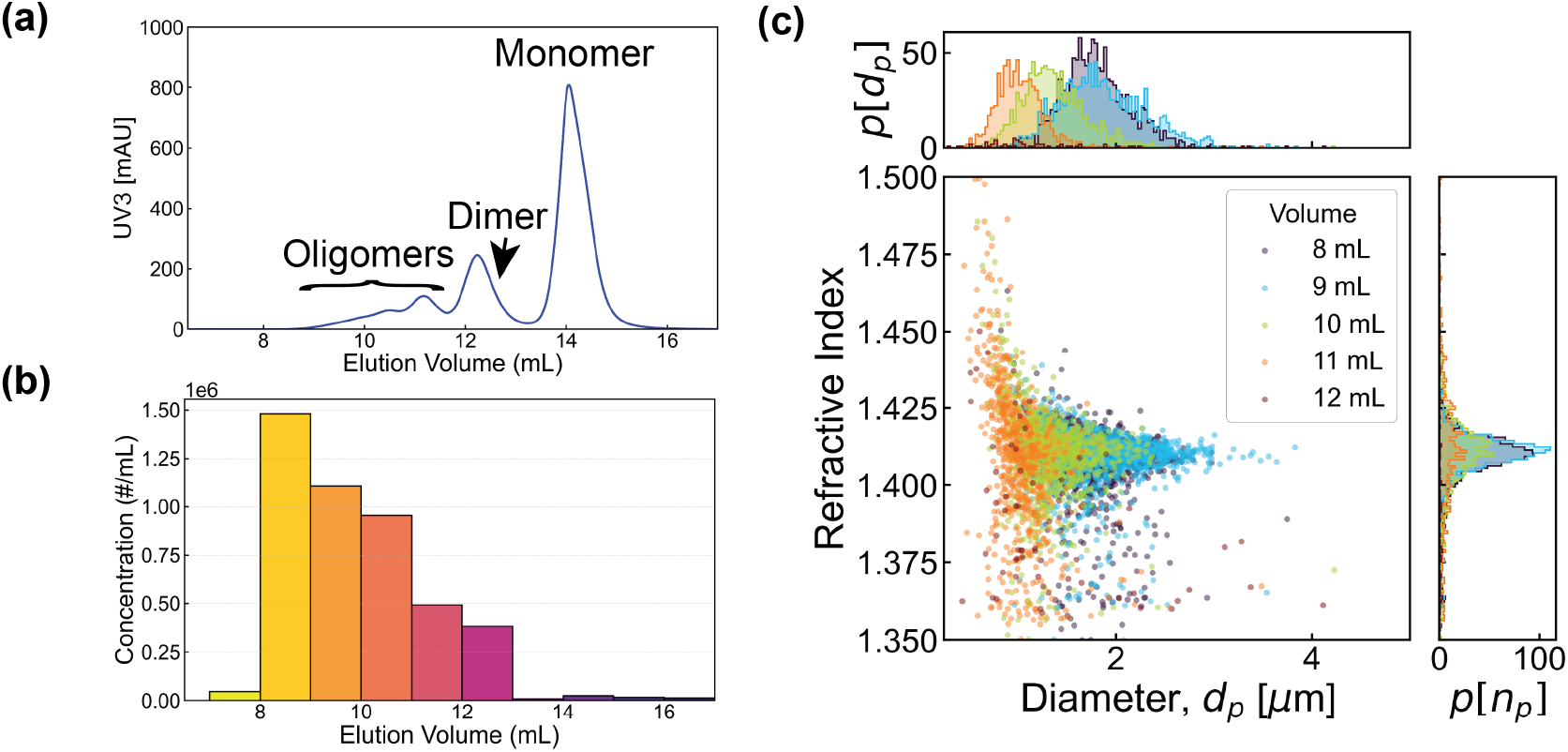
BSA condensate formation is strictly dependent on oligomerization. (a) Size-exclusion chromatography (SEC) UV absorbance (UV3 [mAU]) profile of BSA, showing distinct peaks corresponding to monomers, dimers, and higher-order oligomers based on their elution volume (mL). (b) Total particle concentration (measured by holographic characterization) for collected SEC fractions (07 through 16). The highest particle counts are found in fractions 08-12, corresponding to the higher oligomer and dimer peaks. (c) Two-dimensional scatter plots of refractive index versus particle diameter (*d*_*p*_) for the corresponding SEC fractions (A7-A16). The characteristic dense condensate population, identified by a high refractive index (∼ 1.40–1.42), is present only in the early fractions (A7-A11) that contain the higher-order oligomers. Later fractions (A12-A16), which contain primarily dimers and monomers, are devoid of this high-refractive-index population. These results demonstrate that higher-order oligomerization is a prerequisite for BSA to form biomolecular condensates.

The holographic analysis of these fractions provided a definitive molecular criterion for assembly (Figure 4b-c). Fractions containing predominantly monomers and dimers (elution volumes > 12 mL) failed to form any detectable high-density condensates; they either remained soluble or formed loose, low-index aggregates. In sharp contrast, the characteristic population of high-refractive-index condensates (*n*_*p*_ ≈ 1.40–1.42) appeared *exclusively* in the early elution fractions (7–11 mL) enriched with higher-order oligomers.

This result establishes that oligomerization is not merely a side product of crowding but a strict prerequisite for phase separation in this system. We propose that individual BSA monomers lack sufficient interaction surface area (“stickers”) to support the condensed liquid state. Higher-order oligomerization amplifies the effective valence of the protein complex, enabling it to engage in the cooperative, multivalent interactions necessary to nucleate and sustain a biomolecular condensate.

## Discussion

The results presented here establish an oligomer-driven mechanism for phase separation of a structured globular protein, extending multivalency-based theories of condensate formation beyond intrinsically disordered systems.^4,23,24^ By combining size-exclusion chromatography with holographic characterization, we show that only higher-order BSA oligomers generate dense condensates under conditions where monomers and dimers remain non-condensing or aggregate, indicating that quaternary structure endows a globular protein with emergent multivalency. This conclusion is consistent with recent work demonstrating that oligomerization domains or coiled-coils can lower condensation thresholds and enable phase separation of modules that do not independently condense.^30–32^

Figure 5 summarizes a mechanistic model that integrates our observations of oligomer dependence, pH sensitivity, and crowding thresholds into a unified picture. In this model, higher-order oligomers provide the multivalent scaffold required for phase separation, short-range hydrophobic contacts supply the salt-resistant cohesive energy, and protonation state tunes condensate mesh size by modulating electrostatic repulsion between oligomeric sub-units. The external crowding environment then selects between soluble, aggregated, and condensed states: only when the depletant concentration and polymer size exceed critical values does the system access a stable dense liquid rather than falling into kinetic aggregation traps. This framework positions BSA condensates as a globular-protein counterpart to hydrophobically driven phase separation in disordered systems and rationalizes why condensate density is pH-tunable yet largely insensitive to added salt.^28,29,33–35^

**Figure 5.**
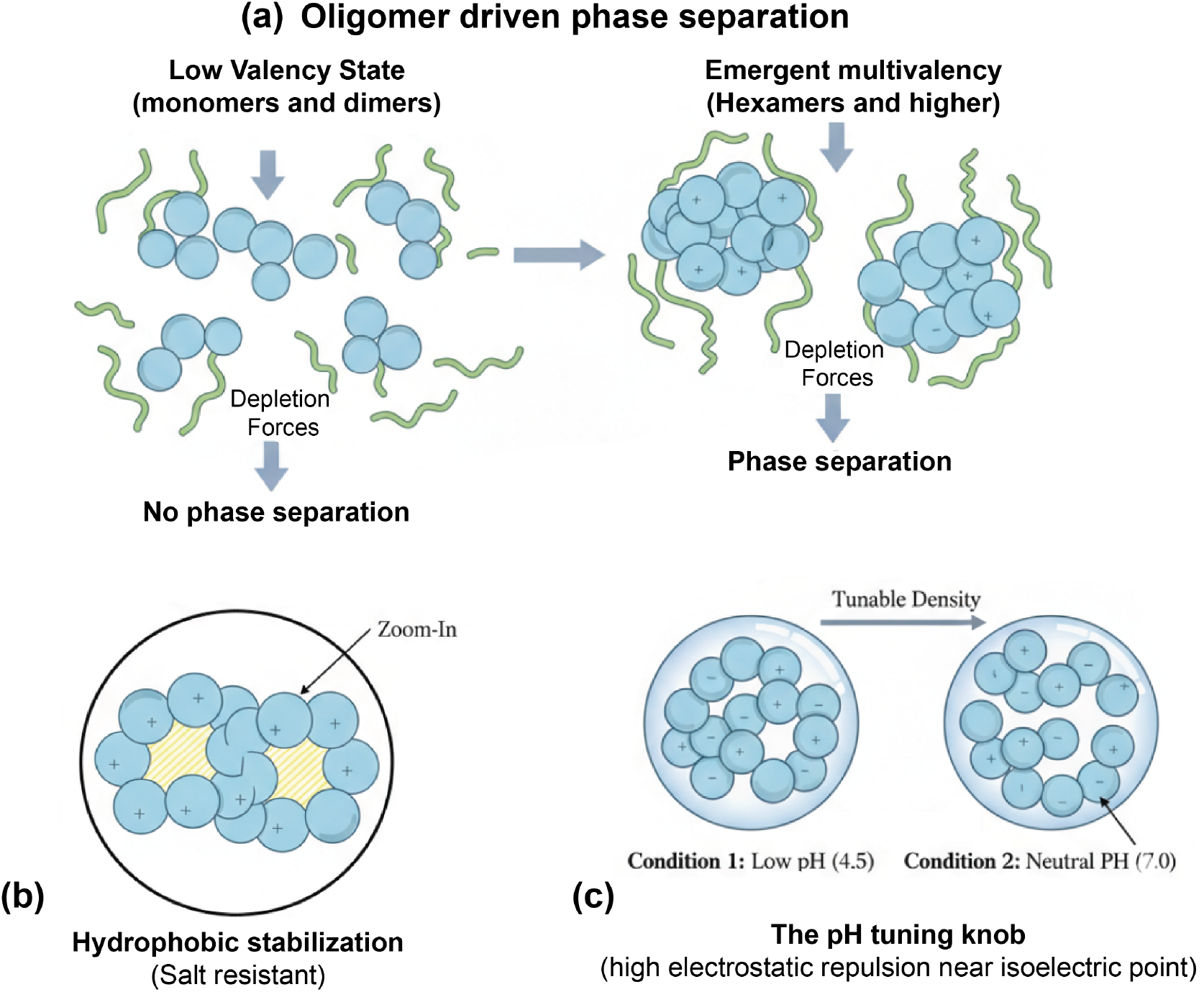
Schematic model for oligomer-driven LLPS of BSA under crowding. Higher-order BSA oligomers provide the multivalent scaffold required for phase separation, whereas monomers and dimers remain non-condensing or form low-index aggregates. Short-range, salt-resistant hydrophobic contacts supply the cohesive energy that stabilizes the dense phase, while protonation state tunes condensate mesh size by modulating electrostatic repulsion between oligomeric subunits. The external crowding environment, set by PEG concentration and molecular weight, selects between soluble, aggregated, and condensed states, with only sufficiently strong and long-ranged depletion attractions yielding stable, high–refractive-index condensates.

A key advance of this work is the use of inline digital holographic microscopy to distinguish, in a label-free and high-throughput manner, dense liquid condensates from amorphous aggregates based on their single-particle optical fingerprints. By extracting both size and refractive index for thousands of individual assemblies, holographic characterization resolves populations that are indistinguishable by bulk turbidity or conventional imaging and directly links oligomeric state to condensate material properties. Beyond basic biophysics, mis-regulated oligomerization and aberrant aggregation underlie numerous aggregation-related pathologies, including amyloid disorders and aggregation of therapeutic proteins.^36,37^ Because quaternary structure often determines whether proteins remain soluble, form reversible condensates, or enter irreversible aggregates, methods that report on both oligomerization and condensate properties—such as the combination of SEC and holographic particle characterization used here—provide a general framework for dissecting these transitions at the molecular level.

More broadly, an oligomer-driven, hydrophobically stabilized mechanism for BSA phase separation suggests that globular proteins can access condensate states without extensive disorder by assembling multivalent scaffolds through oligomerization, complementing models that focus on intrinsically disordered regions and modular interaction domains.^4,23,24^ Practically, the sensitivity of BSA to modest changes in ionic strength, pH, and crowding highlights a narrow energetic balance between reversible condensation and irreversible aggregation, directly relevant for protein stability in therapeutic formulations and crowded physiological fluids.^9,27,36^ Future work coupling holographic characterization with perturbations such as chaotropic agents, hydrophobicity modulators, or sequence variants could further dissect the microscopic interactions that govern globular-protein condensates and test whether oligomer-driven multivalency is a widespread strategy for modulating phase separation.^38–42^ Follow-up studies will be required to determine whether such environmentally tunable phase behaviors are not only biophysically accessible but also crucial for controlling cellular signaling and stress responses, as suggested for bacterial condensates.^43,44^

## Experimental

### Reagents

Bovine serum albumin (BSA, lyophilized powder, protease-free, MW 66,400 g mol^−1^) was purchased from Tocris Biotechne (EC232-936-2, United Kingdom). Sodium chloride (NaCl) was obtained from Fisher Scientific (S671-3). Poly(ethylene glycol) (PEG 8,000) was purchased from Bioworld (Cat.# 705631) and bioPLUS Fine Research Chemicals. Sodium phosphate monobasic (NaH_2_PO_4_) and sodium phosphate dibasic (Na_2_HPO_4_) were obtained from Sigma-Aldrich (Cat.# S9638 and S9763). All solutions were prepared in deionized water produced by a Milli-Q purification system (18.2 MΩ cm at 24 °C).

### Buffer and Sample Preparation

A 1 M sodium phosphate stock at pH 7.0 was prepared by mixing appropriate volumes of 1 M NaH_2_PO_4_ and 1 M Na_2_HPO_4_ and adjusting the pH with NaOH or HCl as needed. Working buffer solutions at 0.10 M, 0.010 M, and 0.0010 M total phosphate concentration were obtained by dilution of the stock with Milli-Q water. Unless otherwise noted, buffers contained 150 mM NaCl.

NaCl stock solutions were prepared by dissolving NaCl in the corresponding phosphate buffer to the desired concentration and filtering through a 0.22 *μ*m membrane prior to use.

### Protein Reconstitution

Lyophilized BSA was dissolved in sodium phosphate buffer (pH 7.2, 100 mM total phosphate, 150 mM NaCl) to a nominal concentration of 750 *μ*M. Protein concentration was determined by UV–vis absorption at 280 nm using a NanoDrop spectrophotometer (DeNovix, DS11) and an extinction coefficient of *ε*_280_ = 44,289 M^−1^ cm^−1^. Samples were stored at 4 °C and used on the same day.

### Size-Exclusion Chromatography of BSA Oligomers

BSA oligomeric states were separated by size-exclusion chromatography (SEC) on a Superdex 200 Increase 10/300 GL column (Cytiva, 28-9909-44) equilibrated with 100 mM sodium phosphate buffer (pH 7.0) containing 150 mM NaCl. The column was operated at mL min^−1^ on a Bio-rad FPLC system (Biorad, NGC) at room temperature. After equilibration with at least three column volumes of buffer, BSA samples (e.g., 0.5–1.0 mL at 100–300 *μ*M) were injected and elution was monitored at 280 nm.

Fractions of 1.0 mL were collected across the elution profile. Oligomeric states (monomer, dimer, higher-order oligomers) were assigned based on elution volumes relative to a calibration curve obtained using standard proteins of known molecular weight. Protein concentrations in each fraction were determined by UV absorption at 280 nm using Beer–Lambert’s law, *c* = *A/*(*εl*), where *c* is the molar concentration, *A* is the absorbance at 280 nm, *l* is the path length (cm), and *ε* is the molar extinction coefficient.

### Condensate Formation

For unfractionated BSA, condensates were formed by dissolving 50.0 mg of BSA in 1.0 mL of sodium phosphate buffer (pH 7.0, 100 mM phosphate, 150 mM NaCl) to yield a stock protein concentration of approximately 750 *μ*M. PEG 8,000 was then added from a concentrated stock solution to achieve the desired final concentration (typically 10–30% w/v), and the mixture was gently mixed until the PEG was fully dissolved. Samples were incubated for 15 min at room temperature and then stored at 4 °C until measurement.

For SEC fractions, 70 *μ*L of each protein fraction was mixed with 80 *μ*L of PEG 8,000 solution in the corresponding sodium phosphate buffer to obtain a final PEG concentration of 20% w/v and 150 mM NaCl at the indicated pH values. Samples were incubated at ambient temperature for [10–30] min prior to holographic characterization.

### Holographic Particle Characterization

Condensate size and refractive index distributions were measured using an inline holographic particle characterization instrument (xSight, Spheryx Inc., USA). In this approach, samples are flowed through a microfluidic channel (xCell, Spheryx Inc., USA) illuminated by a collimated laser beam (*λ* = 450 nm). Each particle scatters the incident light, producing an interference pattern (hologram) with the unscattered beam. Holograms are recorded within a viewing volume of approximately 150 *μ*m × 120 *μ*m × 50 *μ*m using a 40× objective and a digital camera.

For each measurement, 30 *μ*L of sample was loaded into an xCell device and flowed through the channel at a controlled rate set by the instrument software. Recorded holograms were analyzed using the xSight Total Holographic Characterization software, which fits each hologram to Lorenz–Mie scattering theory to extract the particle’s effective diameter, refractive index, and morphological symmetry at the single-particle level. ^45,46^ A typical run lasted ∼15 min, yielding measurements for 10^3^–10^6^ particles in the size range 0.5–10 *μ*m. Particle concentrations were estimated by counting the number of particles within the known sample volume analyzed.

## Acknowledgement

The authors thank Julian von Hofe and Moeka Sasazawa of the Saurabh research group for helpful discussions and inputs. The authors thank Maxwell Cheong for help with data collection. This study was supported by the National Institutes of Health through award 1R35GM157103 to S.S. The xSight instrument used for this study was acquired as shared instrumentation with support from the MRSEC program of the NSF under award DMR-1420073.

